# Resources for Interpreting Variants in Precision Genomic Oncology Applications

**DOI:** 10.1101/144766

**Authors:** Hsinyi Tsang, Durga Addepalli, Sean R. Davis

**Affiliations:** Center for Cancer Research, National Cancer Institute, National Institutes of Health, Bethesda, MD, USA; Center for Biomedical Informatics and Information Technology, National Cancer Institute, National Institutes of Health, Gaithersburg, MD, USA; Attain, LLC, McClean, VA, USA

## Abstract

Precision genomic oncology–applying high throughput sequencing (HTS) at the point-of-care to inform clinical decisions–is a developing precision medicine paradigm that is seeing increasing adoption. Simultaneously, new developments in targeted agents and immunotherapy, when informed by rich genomic characterization, offer potential benefit to a growing subset of patients. Multiple previous studies have commented on methods for identifying both germline and somatic variants. However, interpreting individual variants remains a significant challenge, relying in large part on the integration of observed variants with biological knowledge. A number of data and software resources have been developed to assist in interpreting observed variants, determining their potential clinical actionability, and augmenting them with ancillary information that can inform clinical decisions and even generate new hypotheses for exploration in the laboratory. Here, we review available variant catalogs, variant and functional annotation software and tools, and databases of clinically actionable variants that can be used in an ad hoc approach with research samples or incorporated into a data platform for interpreting and formally reporting clinical results.

## 1 Introduction

Genomic technologies and approaches have transformed cancer research and have led to the production of large-scale cancer genomics compendia (*International Cancer Genome Consortium* n.d.; Cancer Genome Atlas Research Network et al. 2013). The resulting molecular characterization and categorization of individual samples from such compendia has driven development of molecular sub-types cancers as well as enhanced understanding of the molecular etiologies of carcinogenesis (Cancer Genome Atlas Network 2012; Cancer Genome Atlas Research Network 2015; Network 2008). The development of novel and effective targeted therapies has proceeded in parallel with and been accelerated by deeper, faster, and broader genomic characterization (Blumenthal, Mansfield, and Pazdur 2016), enabling early application of molecular characterization at the point of care to inform clinical decision-making (Flaherty et al. 2012; Shaw et al. 2013; Maemondo et al. 2010; Druker et al. 2006) and to address resistance to primary therapy (Ai and Tiu 2014). Genomic characterization also has applications in immune approaches to cancer. For example, chimeric antigen receptor T-cell (CARt) therapy have shown great success in diseases with well-characterized antigens that are relatively tumor-specific (Grupp et al. 2013) as identified by genomic profiling. Variously referred to as preci-sion oncology (Sohal et al. 2015), genomics-driven oncology (Garraway 2013), genomic oncology, and even simply as precision medicine, the paradigm of applying high-throughput genomic approaches to patient samples is rapidly changing the landscape of oncology care and clinical oncology research.

Conventional approaches to clinical trials design may be inadequate due to molecular heterogeneity of tumors derived from a single primary tissue (Simon 2016), leading to the adoption of basket, umbrella, and hybrid trials designs. A number of studies are ongoing to determine feasibility and potential impact of precision genomic oncology at the point-of-care (Cheng et al. 2015; *NCI-MATCH Trial (Molecular Analysis for Therapy Choice)* n.d.; Lopez-Chavez et al. 2015). In addition to studies focused on identifying targetable mutations, immune-based therapeutic approaches are also being informed by HTS applied to patient samples (Bethune and Joglekar 2017; Chalmers et al. 2017; Faltas et al. 2016).

One of the most recent developments in the field of precision oncology is the approval of Pembrolizumab (Keytruda), a anti-PD-1 antibody that functions as a checkpoint inhibitor, by the US Food and Drug Administration for treatment of solid tumors that show genetic evidence of mismatch repair and therefore carry very high mutational burdens (Le et al. 2017). Pembrolizumab was previously approved for use in melanoma, but the most recent approval is the first that is targeting allows a drug to be used in a non-tissue-specific context in patients showing a specific genomic marker in any solid tumor (Garber 2017).

As with any clinical testing modality, whether in a research setting or at the point-of-care, a clear understanding of the goals of applying the test is necessary when first designing the test and its validation. However, the flexibility and number of potential data items that arise from even a limited application of HTS has lead the US Food and Drug Administration (FDA) has begun to define its regulatory role (FDA 2015) and, critically, how existing knowledge bases can be applied in real time to address findings from clinical HTS testing (FDA 2016).

This review aims to provide an organized set of biological knowledge bases with relevance to the interpretation of small variants, defined as single nucleotide variants or short (on the order of 20 base pairs or fewer) insertions and deletions. The catalogs of observed variants section lists large-scale catalogs of variants, useful for filtering known common polymorphisms and identifying previously-identified cancer variants. When a variant observed in a clinical sample has not been seen but appears to affect the protein coding sequence, the functional annotation resources section presents a sampling of some of the most common software and databases for predicting the impact on protein function. Finally, we catalog several data products and knowledgebases have been developed to provide decision support (with strong disclaimers and caveats) directly linking observed variants to clinical intervention in point-of-care HTS applications. Integrating the various data sources described in this review with variants observed in individual patients can be accomplished with combinations of software tools for the manipulation of variant datasets.

### 1.1 Catalogs of observed germline and somatic variants

Databases of observed variation in normal populations, diseased individuals, and cancer compendia form the map onto which observed variants in patients are projected. Because of the vast quantities of genomic data and, specifically, DNA variants, there is a tension between providing rich, highly-curated information about individual variants and producing the largest possible catalog of variants with manageable levels of curation. This section reviews some of the available catalogs (Table 1) of genomic variation observed in the germline as well as those that appear in tumors as somatic mutations. Note that many of the databases mentioned below overlap in data sources (some nearly completely), but they may differ in the amount and depth of curation, additional metadata added to each variant, speed of updates, and methods or formats for access.

**Table 1:**
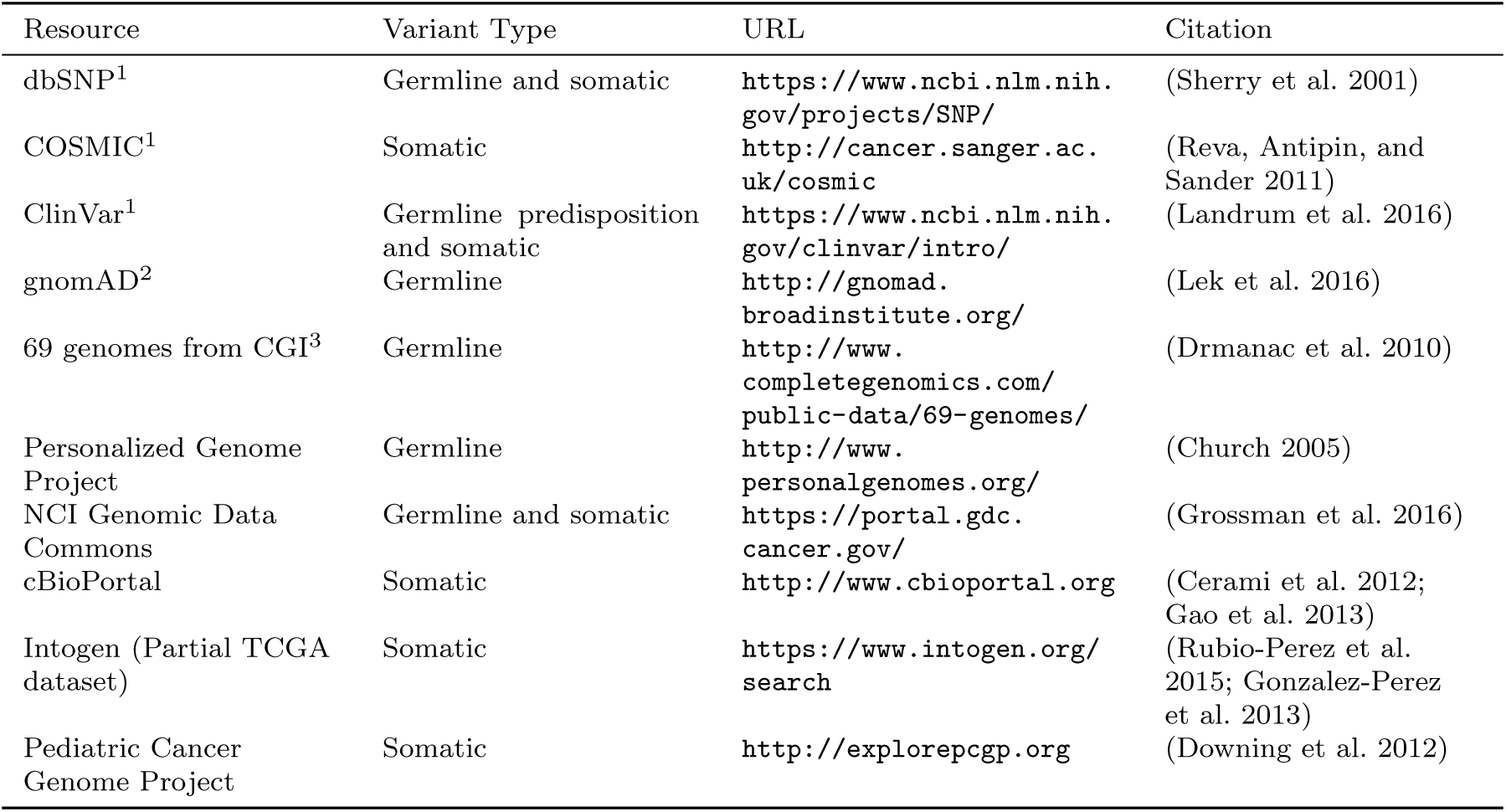
Catalogs of germline and somatic variants. The most commonly used catalogs include dbSNP, COSMIC, ClinVar, and gnomAD. ^1^Primary resources useful for all studies. ^2^Particularly useful for exome sequencing projects. ^3^Useful if the Complete Genomics platform was used.

### 1.2 Germline

Comprehensive catalogs of germline variants inform decisions about the frequency of variants as seen in the general population as well as to identify variants that are annotated as cancer-associated. In the context of tumor sequencing, common variants are unlikely to be genomic drivers of carcinogenesis and are often filtered from a report of potential somatic variants. This filtering process is particularly important when tumor sequencing is not accompanied by matched normal sequencing. Additional germline databases that catalog disease-associated variants can be useful to begin to address familial risk and potentially pharmacogenomic loci (Wheeler et al. 2013; Relling and Evans 2015).

Perhaps the oldest of the variant catalogs, dbSNP contains 325,658,303 individual variant records (build 150, accessed May 30, 2017) and is available in multiple formats, searchable, and linked to records in literature and other data resources and databases. While the vast majority of variants in dbSNP have been observed in individuals without cancer, somatic variants are included and annotated in the database. Because dbSNP is driven by community submission of variants, levels of evidence vary among individual variants. The genome Aggregation Database, or gnomAD, (Lek et al. 2016; *gnomAD browser* n.d.) contains information from 123,136 exomes and 15,496 whole-genomes from unrelated individuals sequenced as part of various disease-specific and population genetic studies (accessed May 30, 2017). These data were collected by numerous collaborations, underwent standard processing, and unified quality control and results area accessible as a search-able online database and as a downloadable VCF-format text file. ClinVar (Landrum et al. 2016), maintained by the NIH National Center for Biotechnology Information (NCBI), is a freely available archive for interpretations of clinical significance of variants for reported conditions. Entries in ClinVar are taken directly from submitters and represent the relationship between variants and clinical significance. When multiple submissions concerning a single variant are available, ClinVar supplies high-level summaries of agreement or disagreement across submitters. Importantly, though, clinical significance in ClinVar is reported as supplied by the submitter. The Personalized Genome Project (Church 2005) provides a limited number of fully open-access genome sequencing results provided by volunteers with trait surveys and even some microbiome surveys of participants. A catalog of germline variants derived from 69 genomes sequenced using the Complete Genomics sequencing platform (Drmanac et al. 2010) may be useful for groups who have data generated from the same platform, particularly for identifying sequencing-platform-specific false positive results.

### 1.3 Somatic

Whereas databases of germline variants are useful to filter out variants unlikely to be directly involved in carcinogenesis, databases of somatic variants are useful to identify variants and their frequencies as observed in tumors. In some cases, identified variants may be associated with specific tumor types, offering mechanistic clues, particularly in the rare cancer setting where biological understanding may be limited.

Several catalogs of somatic variants have, at their core, variants derived from The Cancer Genome Atlas (TCGA). These databases vary in the pipelines used to define the variants, the level of annotation associated with individual variants, the proportion of TCGA included, and methods for accessing or querying. Recently, National Cancer Institute (NCI) has established the Genomic Data Commons (GDC) to harmonize clinical information and genomic results across enterprise cancer datasets (Grossman et al. 2016), particularly those funded by NCI, such as TCGA. In addition to the adult tumors profiled as part of the TCGA, the NCI GDC also contains data from several pediatric tumors profiled as part of the Therapeutically Applicable Research To Generate Effective Treatments (TARGET) project (*Therapeutically Applicable Research to Generate Effective Treatments (TARGET)* n.d.). Cancer cell line data from the Cancer Cell Line Encyclopedia (CCLE) are also included (Barretina et al. 2012) in the GDC data collection. The GDC is a modern data platform that provides multiple access methods including a programmatic application programming interface (API), data file download, and web browser based text and graphical queries and visualization. The International Cancer Genome Consortium (ICGC) is a large, international collaboration with a collection of 76 studies (including TCGA studies) encompassing 21 tissue primary sites. Like the NCI GDC, the ICGC data portal provides modern data platform approaches to data access, visualization, and query (Zhang et al. 2011). The Catalog of Somatic Mutations in Cancer (COSMIC) database is perhaps the largest and best-known cancer variant database. It presents a unified dataset consisting of curated cancer variants for specific genes as well as genomic screens from projects such as TCGA. Several other cancer variant data resources are listed in Table 1.

## 2 Functional Annotation Resources

When faced with variants with little or no literature or database support, differentiating those that variants that are likely to be deleterious, perhaps contributing to carcinogenesis, versus those that likely are tolerated by the cell is a critical task, particularly in the setting of clinical precision genomic oncology. Note that determing that a variant is deleterious is not likely to result in a change in diagnosis, prognosis, or therapy. However, prioritizing variants for further study, research interest, and for discussion in forums such as a molecular tumor board is a valuable and necessary aspect of applying genomic technologies in the clinical arena.

A number of algorithms and methods have been developed to predict the effect of observed variants on protein structure and function as well as the potential for clinical impact. These prediction methods utilize features of the variant and its context such as sequence identity, sequence conservation, evolutionary relationship, protein primary and secondary structure, entropy based protein stability and approaches such as clustering based on sequence alignments and machine learning. Some of them are specific to the type of variant or mutation, some to a disease type, and some more general. Therefore, applying these functional annotational tools and interpreting the results in a clinical or research setting may require significant human curation before being recognized as clinically actionable. Here we present a review of a representative set of approaches for predicting pathogenicity of different variants. For a comprehensive list of prediction tools and their details see Table 2. For more detailed scientific and technical explanations of these methods, we refer the reader to a comprehensive review (Addepalli 2014).

**Table 2:**
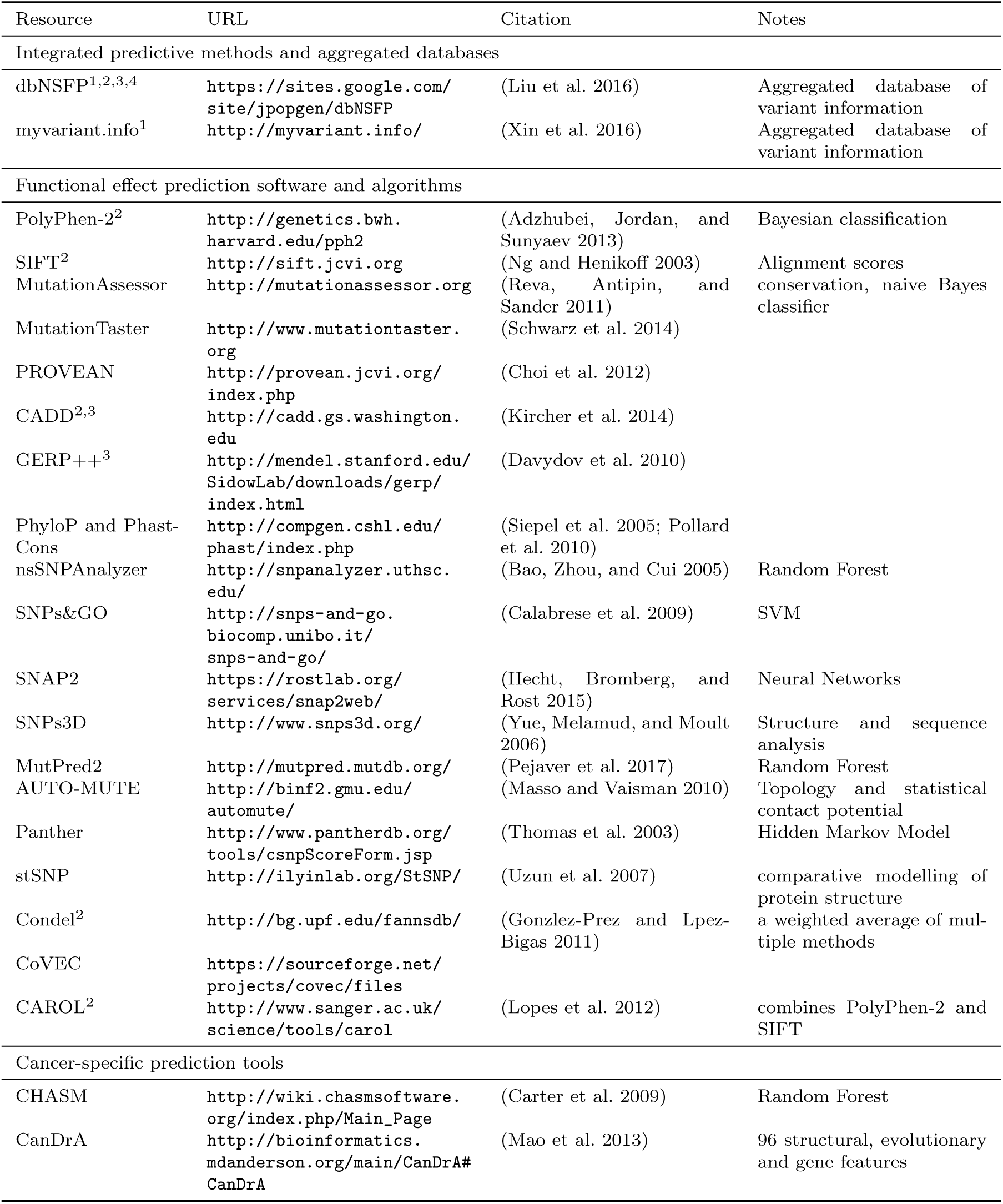
Tools, software, and databases for functional prediction and annotation of variant impact. ^1^Aggregated databases combine outputs of other databases and algorithms are, therefore, effcient resources to use in annotation pipelines. Adding these resources to observed variants is supported software in table 4 including Ensembl VEP software (noted^2^ in this table), Annovar (noted^3^), and snpEff (noted^4^).

### 2.1 SIFT

Sorting Intolerant From Tolerant, or SIFT, predicts functional impacts of amino acid substitutions (Ng and Henikoff 2003) is one of the earliest variant effect prediction tools and represents the class of prediction algorithms that utilizes protein conservation. It has since been updated and an online version of the tool is available (Kumar, Henikoff, and Ng 2009). SIFT uses sequence homology, as measured by protein-level conservation, to classify variants based as tolerated or deleterious based on the associated protein coding changes. SIFT has served as a benchmark against which other methods are compared because of its relative simplicity. SIFT considers the type of amino acid change induced by a genomic variant and the position at which the change/mutation occurs. SIFT relies on the presence of sequences from which conservation can be determined; variants for which such databases are limited will potentially lack robust predictions.

### 2.2 PolyPhen-2

Polymorphism Phenotyping v2, or PolyPhen2, predicts the effecting of coding nonsynonymous SNPs on protein structure and function and annotates them (Adzhubei, Jordan, and Sunyaev 2013). This algorithm uses a naive Bayes approach to combine information across a panel of 3D structural, sequence-based, and conservation-based features. Trained on two datasets, HumDiv and HumVar, and associated non-deleterious controls, the PolyPhen2 algorithm represents a class of multivariate prediction algorithms that employ machine learning and multiple features of variant impact.

### 2.3 Mutation Assessor

Mutation Assessor is an algorithm and tool that, like SIFT, uses a conservation-based approach. However, Mutation Assessor also incorporates evolutionary information in an attempt to account for shifts in function between subfamilies of proteins (Reva, Antipin, and Sander 2011), potentially extending the functional annotation of variants to “switch of function” as well as loss or gain of function. By quantifying the impact to conserved residues both globally and within subfamilies (residues that distinguish subfamilies from each other are thought to be less tolerant to change), Mutation Assessor defines a functional impact score to predict which variants are likely to be deleterious.

### 2.4 CONDEL

The CONsensus DELeteriousness, or CONDEL score, is an integrated prediction method for missense mutations that is relatively easy to extend with additional prediction resources (Gonzlez-Prez and Lpez-Bigas 2011). Originally implemented as a weighted average of the normalized scores from the output of two computational tools, Mutation Assessor and FATHMM, CONDEL can be extended or adapted to data at hand and represents an “aggregator” approach to variant effect prediction. Condel scores can be derived for a limited set of specified mutations via an online web application. The Ensembl database provides a variation of position-specific CONDEL predictions that combine SIFT and Polyphen-2 for every possible amino acid substitution in all human proteins.

### 2.5 CHASM

Cancer-specific High-throughput Annotation of Somatic Mutations, or CHASM, is a computational method that identifies and prioritizes the missense mutations likely to enhance tumor cell proliferation (Carter et al. 2009). CHASM uses machine learning to classify putative “driver” cancer mutations as distinct from “passenger” mutations. Training the CHASM model employed in-silico simulation to generate realistic “passenger” mutations, specifically modeled to represent variant context and genes that are observed in cancer settings. Multiple features of the variants including their DNA and protein contexts were then used to build a machine learning approach that attempted to maximize the specificity of separating driver mutations from passenger mutations. CHASM represents a relatively specific algorithm focused not on “deleteriousness” but, rather, on the likelihood that an observed variant is a cancer “driver”.

### 2.6 dbNSFP

Recognizing that applying all of the effect prediction tools available is potentially challenging, Liu et al. 2016 developed a database that aggregates predictions for *all* possible SNVs associated with coding changes (in Gencode gene models). With more than ten different prediction algorithms and extensive additional annotation, this database can be a useful one-stop-shop for adding annotations to variant datasets. The snpEff suite (described below) can be used in conjunction with dbNSFP to effciently annotate SNPs with the potential to effect coding genes.

## 3 Clinical Actionability

The ultimate goal for many of the above-mentioned resources is to develop an individualized approach to the diagnosis, prevention and treatment of cancer, or precision oncology. However, despite recent advances in HTS, determining the clinical relevance of experimentally observed cancer variants remains a challenge in the application of HTS in clinical practice. Difficulties in differentiating driver and passenger mutations, lack of standards and guidelines in reporting and interpretation of genomic variants, lack of clinical evidence in associating genomic variants to clinical outcome, lack of resources to disseminate clinical knowledge to the cancer community and the precise definition of actionability have been reported to contribute to the bottleneck (M. M. Li et al. 2017; Prawira et al. 2017; Uzilov et al. 2016; Hedley Carr et al. 2016). Comprehensive resources linking experimentally determined cancer variants and clinical actionability have been developed to address some of these challenges and address various aspects of translating research results into clinical valuable information to support clinical decisions in precision oncology (See Table 3). In recognition of the fact that central curation of information regarding actionability is extremely challenging, several of the resources below use crowdsourcing as a means of gathering updates and enhancing curation efforts. In addition to a web interface, some tools provide additional access via API, mobile app, and/or social media tagging to facilitate dissemination of information and enhance accessibility. While some of these tools share similar functions, in the section below, we highlight distinct features and capabilities for a representative set of resources that might be used as a “starter” set for clinical annotation of variants.

**Table 3:**
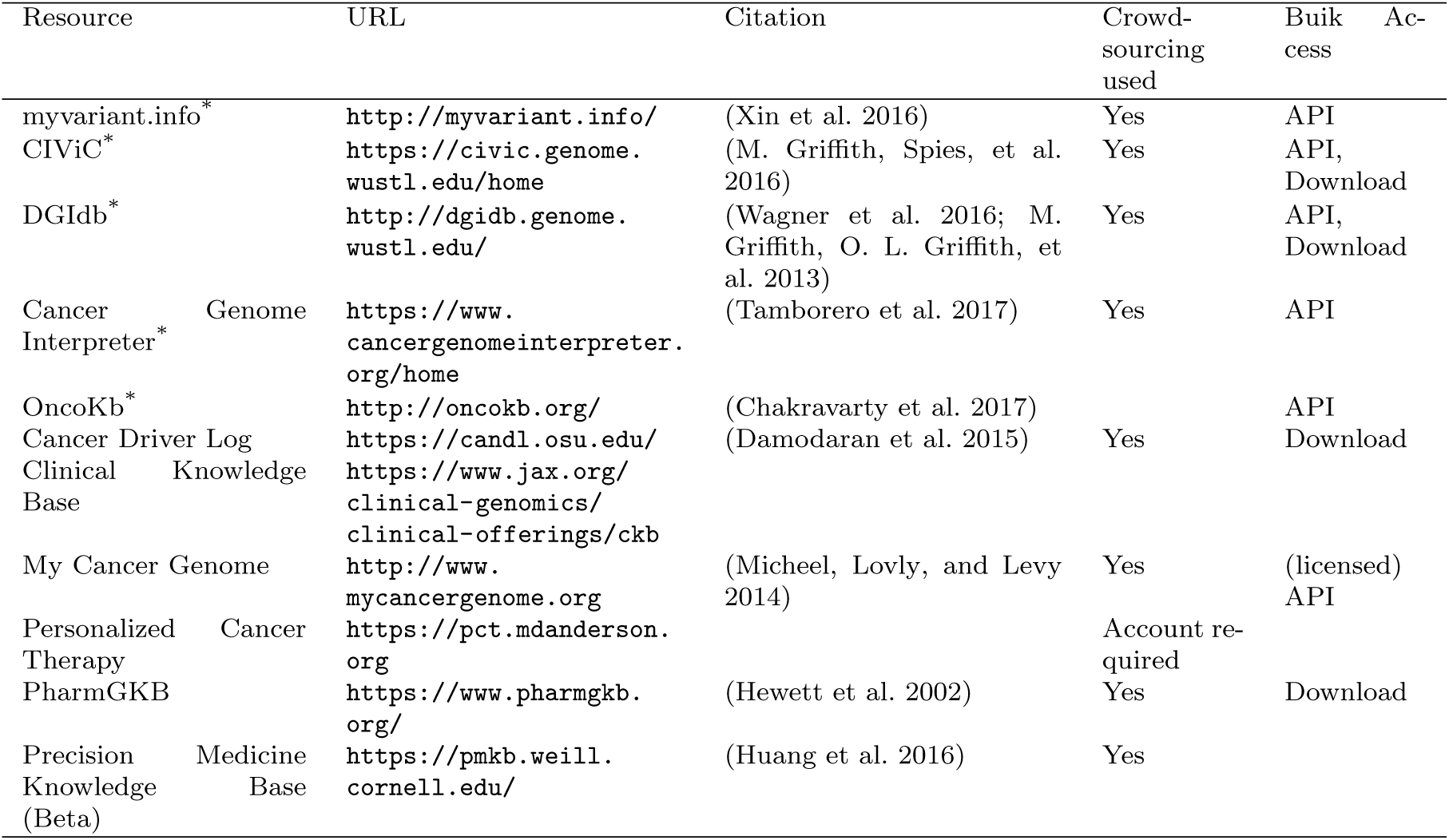
In a clinical setting, these databases are the most relevant, as they are maintained to provide clinically actionable and curated content. While evalutation of each database by both clinical and informatics team members, databases marked with “*” are maintained, recently (or continuously) updated, and curated. The myvariant.info database includes both CiVIC and Cancer Genome Interpreter data. The last column in the table notes bulk access approaches as these are relevant when including databases in an annotation pipeline or automated report.

The myvariant.info database is one of the newest and attempts to provide a “one-stop-shop” for variants. It is included in this section because it has recently incorporated the CIViC and Cancer Genome Interpreter databases. In addition, it provides annotations for SNVs from multiple other data sources (a growing list, so see the site for updates) and aggregates functional annotations for variants present in its database, making it a good all-around tool for cancer variant annotation. It is available as a performant web API only at this time.

Clinical Interpretation of Variants in Cancer (CIViC) is an open access and open source platform for community-driven curation and interpretation of cancer variants. It is based on a crowdsourcing model where individuals in the community can contribute to produce a centralized knowledge base with the goal of disseminating knowledge and encouraging active discussion. Users, including patients, patient advocates, clinicians and researchers, can participate, along with community editors, in various stages of interpreting the clinical significance of cancer variants using standards and guidelines developed by community experts (M. M. Li et al. 2017; M. Griffith, Spies, et al. 2016).

The Drug Gene Interaction Database (DGIdb) is an open source and open access platform for gene and drug annotation for known interaction and potential druggability. Users can can cross-reference genes of interest and drugs against up to 15 sources and in functionally classified gene categories (Wagner et al. 2016; M. Griffith, O. L. Griffith, et al. 2013). Cancer Genome Interpreter (CGI) identifies mutational events that are biomarkers of drug response or interact with known chemical compounds (Tamborero et al. 2017). PharmGKB is a pharmacogenomic resource for building clinical implementation and interpretation based on annotating, integrating and aggregating knowledge extracted from research-level publications. It provides scored clinical annotation, prescription annotation (drug dosing, prescribing information), as well as pharmacokinetics/pharmacodynamics (PK/PD) annotation, with primary literature reference.

OncoKb contains information on the clinical implication of specific genetic alterations in cancer. Each variant is annotation from multiple sources and scored using Levels of Evidence ranging from Level 1, which includes FDA approved biomarker predictive of response to an FDA-approved drug, to Level 2, which includes variants for which an FDA-approved or standard of care treatment is available, Level 3 and Level 4 contain variants with investigational and hypothetical therapeutic implications, respectively. A similarly structured scoring system is available for indicating therapeutic implications for variants associated with resistance (Chakravarty et al. 2017). Cancer Driver Log (CanDL), an expert-curated database for potential driver mutations in cancer, employs a similar four-level scoring system based on FDA approval, clinical, pre-clinical and experimental functional evidence (Damodaran et al. 2015).

MyCancerGenome (MCG) is a knowledge resource highlighting the implication of tumor mutation on cancer care. It allows users to access its content via a mobile app and provide patient-focused information. Patients can access a database entitled DNA-mutation Inventory to Refine and Enhance Cancer Treatment (DIRECT) for Epidermal Growth Factor Receptor (EGFR) mutation for non-small cell lung cancer (NSCLC). Personalized Cancer Therapy (PCT) at the MD Anderson Cancer Center is a resource for clinical response associated with cancer variants and aims to facilitate patient involvement in biomarker-related clinical trials. Drug effectiveness is associated with a specific biomarker and scored based on prospective clinical study as well as Food and Drug Administration (FDA) approval.

## 4 Tools for manipulating variant datasets

Processing sequence data with the goal of determining variants (somatic or germline) often ends with a file in Variant Call Format (VCF format), a loose, self-describing data standard describing variants along a genome, associated statistical and numeric metrics for each variant, and information integrated from data resources such as those described in the preceding sections (Danecek et al. 2011). An ecosystem of tools, listed in Table 4, has been developed for basic transformations, manipulations, merge operations, and for adding transcript, protein, and higher-level functional annotations to variants in a VCF file. The vt and bcftools software suites perform operations such as slicing by genomic coordinate, data compression, and, importantly variant normalization, rendering variants more readily comparable across resources. Annovar (Yang and K. Wang 2015; K. Wang, M. Li, and Hakonarson 2010) and the SnpEff suite (Cingolani 2012) add annotations relative to gene annotations, including information about transcript and protein-coding changes. The Ensembl Variant Effect Predictor (VEP) utilizes Ensembl gene models to annotate variants in gene context and offers an interesting plugin architecture that supports adding variant information from resources in table **??** (McLaren et al. 2016). Recently, several software developers of variant annotation tools have developed a standard for reporting gene-centric annotations that has simplified post-processing of variants after annotation. Finally, tools such as Vcfanno (Pedersen, Layer, and Quinlan 2016) have been developed that can flexibly add fields to variants in a VCF file based on relatively sophisticated logic and data transformations, reducing the number of tools required to bring a new data resource into the annotation pipeline.

**Table 4:** Software tools for manipulating and adding annotations to variant datasets. Variant calling produces a list of observed variants. The tools in this table are useful for adding biological interpretation and for annotating the variants with information from resources in tables 1, 2, and 3.

| Software | URL | Citation |
| --- | --- | --- |
| vt | http://genome.sph.umich.edu/wiki/Vt | (Tan, Abecasis, and Kang 2015) |
| bcftools | http://www.htslib.org/download/ | (H. Li et al. 2009) |
| ANNOVAR | http://annovar.openbioinformatics.org/en/latest/ | (K. Wang, M. Li, and Hakonarson 2010) |
| Ensembl Variant Effect Predictor (VEP) | http://www.ensembl.org/vep | (McLaren et al. 2016) |
| Snpeff | http://snpeff.sourceforge.net/ | (Cingolani 2012) |
| Oncotator | https://portals.broadinstitute.org/oncotator/ | (Ramos et al. 2015) |
| vcfanno | https://github.com/brentp/vcfanno | (Pedersen, Layer, and Quinlan 2016) |

## 5 Discussion

### 5.1 Pragmatic details

Despite advanced toolsets for manipulating variant files and increasing adoption available standard formats, practical pitfalls and challenges remain to the basic manipulation of variant datasets. Some data resources are available in multiple formats and not all formats contain identical information. Matching variants between resources and observed variants can be challenging, as some variants can be represented validly in multiple forms. Ideally, variants are cataloged with clarity with respect to a reference genome and, whenever possible, using HGVS nomenclature (Dunnen et al. 2016). In spite of increasing awareness and uptake of HGVS standard nomenclature, the critical step of matching variants across tools and databases in assessing clinical significance is still hampered by inconsistencies across tools and databases (Yen et al. 2017). Particularly when handling clinical samples, an information system that provides results from multiple resources when assessing novel variants, incorporates in silico controls when adding or updating data resources (to avoid introducing errors), and adheres to HGVS nomenclature wherever possible in data processing pipelines can increase the likelihood of discovering potentially relevant variants.

### 5.2 Where to start?

This review is meant to be comprehensive, so the reader might wonder “Where do we start?”. While it is difficult to make hard-and-fast recommendations about what resources, tools, and databases are “the best” given the lack of gold-standard datasets on which to base such evalutations, annotations in tables 1, 2, and 3 are meant to provide context for prioritization. The context for sequencing (clinical or not, targeted mutations, trial setting, or novel variant and biomarker discovery) will also drive annotation pipeline development. Not all data resources need to be added simultaneously if developing a pipeline for annotating cancer variants for precision oncology applications. In a clinical setting, targeting the reporting workflow and working with clinicians to understand the most relevant annotations is the most efficient approach to determining relevant resources for annotation. Developing a modular informatics pipeline, perhaps using a computational workflow framework (<u>https://github.com/pditommaso/awesome-pipeline</u>) that can be easily extended and re-run on previously annotated data is helpful to keep pace with the rapidly changing and growing collection of annotation resources. Newer aggregation resources such as myvariant.info offer a wholistic solution (annotation, catalog, and clinical actionability), but with some risk of “lossiness” with respect to the primary resources contained within.

Finally, given the rapid pace of new development in this space, we have established a crowd-sourced list of cancer variant resources for precision medicine available at https://github.com/seandavi/awesome-cancer-variant-databases.

### 5.3 Conclusion

Robust sequencing technologies and increasingly reliable bioinformatics pipelines, combined with parallel development of therapeutics and diagnostics has bolstered the field of precision genomic oncology. However, the sheer number of resources available that can inform the interpretation of small variants is staggering, except for the very few variants with well-established clinical relevance or an associated targeted therapy. This review has highlighted a number of important data resources individually. For other variants, data integration remains a significant hurdle to the rapid turnaround required to apply HTS in a clinical context. Expert panel review (the molecular tumor board) has been effective for some groups (Knepper et al. 2017; Beltran et al. 2015; Sohal et al. 2015) while other groups have adopted a protocol-based approach (Ghazani et al. 2017). Even when molecularly targetable lesions are identified, barriers to delivering therapy have been observed, limiting the impact of precision genomic oncology in some settings (Bryce et al. 2017). Not covered in this review is the increasing utility of HTS in the burgeoning field of immunotherapy, where early efforts to predict response based on HTS results have been promising (R.-F. Wang and H. Y. Wang 2017; Yarchoan et al. 2017; Bethune and Joglekar 2017).

Some interesting trends are evident in the databases and resources presented in this review that highlight the overarching trends in delivering precision medicine. First is the sheer volume and rapid growth of numbers of observations to learn about the spectrum of variation cancer and normal genomes. Projects like GnomAD, COSMIC, and other data sharing efforts enhance precision by cataloging rare variants as well as precise estimates of the frequencies of common variants. Second is the use of crowd-sourcing to produce rich clinical annotation (eg., CiVIC) in response to the need for intensive human interaction to interpret the clinical impact of a variant or its relationship to potential medical intervention. On the other hand, with volumes of data ever-increasing, machine learning techniques drive many of the most commonly-used approaches for assigning scores for impact of observed variants. As well-annotated datasets and variant catalogs grow, application of machine learning will become both more common and more powerful.

While significant progress has been made in applying technology to precision oncology, cancer arises in an individual after a typically complex and incompletely understood set of oncogenic events that are increasingly observable at the molecular level. Progress in cancer prevention, early detection, diagnosis, prognosis, and treatment is increasingly driven by insight gained through the analysis and interpretation of large genomic, proteomic, and pharmacological knowledge bases. Reductionist approaches to cancer biology can achieve only limited success in understanding cancer biology and improving therapy. Cancer is a disease associated with disruption of normal cellular circuitry and processes that leads to abnormal or uncontrolled proliferative growth, characterized by a complex spectrum of biochemical alterations that affects biological processes at multiple scales from the molecular activity and cellular homeostasis to intercellular and inter-tissue signaling. The cancer research community has made great strides in measuring the oncogenic events that lead to the development of cancer and therapy resistance. Because of the complexity inherent in protein networks, intercellular signaling, cellular heterogeneity, and the dynamic nature of cancer, future progress will require a more wholistic approach to precision oncology including multiscale systems and modeling approaches that address the interrelatedness of the biological processes underlying cancer.

## 7 Acknowledgments

This work was supported by the National Cancer Institute Center for Biomedical Informatics and Information Technology and the National Cancer Institute Center for Cancer Research in the Intramural Research Program at the National Institutes of Health.

## References

[1] *International Cancer Genome Consortium*. http://icgc.org. Accessed: 2017-5-15.

[2] Cancer Genome Atlas Research Network et al. “The Cancer Genome Atlas Pan-Cancer analysis project”. en. In: Nat. Genet. 45.10 (Oct. 2013), pp. 1113–1120.

[3] Cancer Genome Atlas Network. “Comprehensive molecular portraits of human breast tumours”. en. In: Nature 490.7418 (Apr. 2012), pp. 61–70.

[4] Cancer Genome Atlas Research Network. “The Molecular Taxonomy of Primary Prostate Cancer”. en. In: Cell 163.4 (May 2015), pp. 1011–1025.

[5] The Cancer Genome Atlas Research Network. “Comprehensive genomic characterization de nes human glioblastoma genes and core pathways”. In: Nature 455.7216 (23 10 2008), pp. 1061–1068.

[6] Gideon M Blumenthal, Elizabeth Mans eld, and Richard Pazdur. “Next-Generation Sequencing in Oncology in the Era of Precision Medicine”. en. In: JAMA Oncol 2.1 (Jan. 2016), pp. 13–14.

[7] Keith T Flaherty et al. “Combined BRAF and MEK inhibition in melanoma with BRAF V600 mutations”. en. In: N. Engl. J. Med. 367.18 (Jan. 2012), pp. 1694–1703.

[8] Alice T Shaw et al. “Crizotinib versus chemotherapy in advanced ALK-positive lung cancer”. en. In: N. Engl. J. Med. 368.25 (20 06 2013), pp. 2385–2394.

[9] Makoto Maemondo et al. “Gefitinib or Chemotherapy for Non-Small-Cell Lung Cancer with Mutated EGFR”. In: N. Engl. J. Med. 362.25 (24 06 2010), pp. 2380–2388.

[10] B J Druker et al. “Long-term benefits of imatinib (IM) for patients newly diagnosed with chronic myelogenous leukemia in chronic phase (CML-CP): the 5-year update from the IRIS study”. In: J. Clin. Oncol. 24.90180 (2006), pp. 6506–6506.

[11] Jing Ai and Ramon V Tiu. “Practical management of patients with chronic myeloid leukemia who develop tyrosine kinase inhibitor-resistant BCR-ABL1 mutations”. en. In: Ther. Adv. Hematol. 5.4 (Aug. 2014), pp. 107–120.

[12] Stephan A Grupp et al. “Chimeric Antigen Receptor-Modified T Cells for Acute Lymphoid Leukemia”. In: N. Engl. J. Med. 368.16 (25 03 2013), pp. 1509–1518.

[13] Davendra P S Sohal et al. “Prospective Clinical Study of Precision Oncology in Solid Tumors”. en. In: J. Natl. Cancer Inst. 108.3 (Sept. 2015).

[14] Levi A Garraway. “Genomics-Driven Oncology: Framework for an Emerging Paradigm”. In: J. Clin. Orthod. 31.15 (20 05 2013), pp. 1806–1814.

[15] Richard Simon. “Genomic Alteration-Driven Clinical Trial Designs in Oncology”. en. In: Ann. Intern. Med. 165.4 (16 08 2016), pp. 270–278.

[16] Donavan T Cheng et al. “Memorial Sloan Kettering-Integrated Mutation Pro ling of Actionable Cancer Targets (MSK-IMPACT): A Hybridization Capture-Based Next-Generation Sequencing Clinical Assay for Solid Tumor Molecular Oncology”. In: J. Mol. Diagn. 17.3 (May 2015), pp. 251–264.

[17] *NCI-MATCH Trial* (Molecular Analysis for Therapy Choice). https://www.cancer.gov/about-cancer/treatment/clinical-trials/nci-supported/nci-match. Accessed: 2017-5-31.

[18] Ariel Lopez-Chavez et al. “Molecular pro ling and targeted therapy for advanced thoracic malignancies: a biomarker-derived, multiarm, multihistology phase II basket trial”. en. In: J. Clin. Oncol. 33.9 (20 03 2015), pp. 1000–1007.

[19] Michael T Bethune and Alok V Joglekar. “Personalized T cell-mediated cancer immunotherapy: progress and challenges”. en. In: Current opinion in biotechnology 48 (Aug. 2017), pp. 142–152. issn: 0958-1669, 1879-0429. doi: 10.1016/j.copbio.2017.03.024. url: http://dx.doi.org/10.1016/j.copbio.2017.03.024.

[20] Zachary R Chalmers et al. “Analysis of 100,000 human cancer genomes reveals the landscape of tumor mutational burden”. en. In: Genome Med. 9.1 (19 04 2017), p. 34.

[21] Bishoy Faltas et al. “Generating a neoantigen map of advanced urothelial carcinoma by whole exome sequencing”. In: J. Clin. Oncol. 34.2 suppl (Oct. 2016), p. 354.

[22] Dung T Le et al. “Mismatch-repair deficiency predicts response of solid tumors to PD-1 blockade”. en. In: Science (Aug. 2017). issn: 0036-8075, 1095–9203. doi: 10.1126/science. aan6733. url: http://dx.doi.org/10.1126/science.aan6733.

[23] Ken Garber. “In a major shift, cancer drugs go ‘tissue-agnostic’”. en. In: Science 356.6343 (16 06 2017), pp. 1111–1112. issn: 0036-8075, 1095-9203. doi: 10.1126/science.356.6343.1111.aturl: http://dx.doi.org/10.1126/science.356.6343.1111.

[24] FDA. Optimizing FDA’s Regulatory Oversight of Next Generation Sequencing Diagnostic Tests Preliminary Discussion Paper. 20 02 2015.

[25] FDA. Draft Guidance for Stakeholders and Food and Drug Administration Sta?. (Aug. 2016).

[26] Heather E Wheeler et al. “Cancer pharmacogenomics: strategies and challenges”. en. In: Nat. Rev. Genet. 14.1 (Jan. 2013), pp. 23–34.

[27] Mary V Relling and William E Evans. “Pharmacogenomics in the clinic”. en. In: Nature 526.7573 (15 10 2015), pp. 343–350.

[28] Monkol Lek et al. “Analysis of protein-coding genetic variation in 60,706 humans”. en. In: Nature 536.7616 (18 08 2016), pp. 285–291.

[29] *gnomAD browser*. http://gnomad.broadinstitute.org/. Accessed: 2017-5-31.

[30] Melissa J Landrum et al. “ClinVar: public archive of interpretations of clinically relevant variants”. en. In: Nucleic Acids Res. 44.D1 (Apr. 2016), pp. D862–8.

[31] G M Church. “The personal genome project”. en. In: Mol. Syst. Biol. 1 (13 12 2005), p. 2005.0030.

[32] Radoje Drmanac et al. “Human genome sequencing using unchained base reads on self-assembling DNA nanoarrays”. en. In: Science 327.5961 (Jan. 2010), pp. 78–81.

[33] Robert L Grossman et al. “Toward a Shared Vision for Cancer Genomic Data”. In: N. Engl. J. Med. 375.12 (22 09 2016), pp. 1109–1112.

[34] *Therapeutically Applicable Research* to Generate Effective Treatments (TARGET). https://ocg.cancer.gov/programs/target. Accessed: 2017-5-31.

[35] Jordi Barretina et al. “The Cancer Cell Line Encyclopedia enables predictive modelling of anticancer drug sensitivity”. In: Nature 483.7391 (29 03 2012), pp. 603–607.

[36] Junjun Zhang et al. “International Cancer Genome Consortium Data Portal a one-stop shop for cancer genomics data”. In: Database 2011 (Jan. 2011).

[37] Kanakadurga Addepalli. “Models Predicting Effects of Missense Mutations in Oncogenesis”. PhD thesis. George Mason University, 2014.

[38] Pauline C Ng and Steven Heniko?. “SIFT: Predicting amino acid changes that affect protein function”. en. In: Nucleic Acids Res. 31.13 (Jan. 2003), pp. 3812–3814.

[39] Prateek Kumar, Steven Henikõ, and Pauline C Ng. “Predicting the effects of coding non-synonymous variants on protein function using the SIFT algorithm”. en. In: Nature protocols 4.7 (25 06 2009), pp. 1073–1081. issn: 1754-2189, 1750-2799. doi: 10.1038/nprot.2009.86. url: http://dx.doi.org/10.1038/nprot.2009.86.

[40] Ivan Adzhubei, Daniel M Jordan, and Shamil R Sunyaev. “Predicting functional effect of human missense mutations using PolyPhen-2”. en. In: Curr. Protoc. Hum. Genet. Chapter 7 (Jan. 2013), Unit7.20.

[41] Boris Reva, Yevgeniy Antipin, and Chris Sander. “Predicting the functional impact of protein mutations: application to cancer genomics”. en. In: Nucleic Acids Res. 39.17 (Jan. 2011), e118.

[42] Abel Gonzlez-Prez and Nuria Lpez-Bigas. “Improving the assessment of the outcome of non-synonymous SNVs with a consensus deleteriousness score, Condel”. en. In: Am. J. Hum. Genet. 88.4 (Aug. 2011), pp. 440–449.

[43] Hannah Carter et al. “Cancer-specific high-throughput annotation of somatic mutations: computational prediction of driver missense mutations”. en. In: Cancer Res. 69.16 (15 08 2009), pp. 6660–6667.

[44] Xiaoming Liu et al. “dbNSFP v3.0: A One-Stop Database of Functional Predictions and Annotations for Human Nonsynonymous and Splice-Site SNVs”. en. In: Human mutation 37.3 (Mar. 2016), pp. 235–241. issn: 1059-7794, 1098-1004. doi: 10.1002/humu.22932. url: http://dx.doi.org/10.1002/humu.22932.

[45] Marilyn M Li et al. “Standards and Guidelines for the Interpretation and Reporting of Sequence Variants in Cancer”. en. In: J. Mol. Diagn. 19.1 (Jan. 2017), pp. 4–23.

[46] A Prawira et al. “Data resources for the identi cation and interpretation of actionable mutations by clinicians”. en. In: Ann. Oncol. 28.5 (Jan. 2017), pp. 946–957.

[47] Andrew V Uzilov et al. “Development and clinical application of an integrative genomic approach to personalized cancer therapy”. en. In: Genome Med. 8.1 (Jan. 2016), p. 62.

[48] T Hedley Carr et al. “Defining actionable mutations for oncology therapeutic development”. en. In: Nat. Rev. Cancer 16.5 (26 04 2016), pp. 319–329.

[49] Malachi Griffith, Nicholas C Spies, et al. “CIViC: A knowledgebase for expert-crowdsourcing the clinical interpretation of variants in cancer”. en. (Jan. 2016).

[50] Alex H Wagner et al. “DGIdb 2.0: mining clinically relevant drug-gene interactions”. en. In: Nucleic Acids Res. 44.D1 (Apr. 2016), pp. D1036–44.

[51] Malachi Grifith, Obi L Grifith, et al. “DGIdb: mining the druggable genome”. en. In: Nat. Methods 10.12 (Dec. 2013), pp. 1209–1210.

[52] David Tamborero et al. “Cancer Genome Interpreter Annotates The Biological And Clinical Relevance Of Tumor Alterations”. en. 20 05 2017.

[53] Debyani Chakravarty et al. “OncoKB: A Precision Oncology Knowledge Base”. In: JCO Precision Oncology 1 (Jan. 2017), pp. 1–16.

[54] Senthilkumar Damodaran et al. “Cancer Driver Log (CanDL): Catalog of Potentially Actionable Cancer Mutations”. en. In: J. Mol. Diagn. 17.5 (Sept. 2015), pp. 554–559.

[55] Petr Danecek et al. “The variant call format and VCFtools”. en. In: Bioinformatics 27.15 (Jan. 2011), pp. 2156–2158.

[56] Hui Yang and Kai Wang. “Genomic variant annotation and prioritization with ANNOVAR and wANNOVAR”. en. In: Nat. Protoc. 10.10 (Oct. 2015), pp. 1556–1566.

[57] Kai Wang, Mingyao Li, and Hakon Hakonarson. “ANNOVAR: functional annotation of genetic variants from high-throughput sequencing data”. en. In: Nucleic Acids Res. 38.16 (Sept. 2010), e164.

[58] Pablo Cingolani. snpE?: Variant effect prediction. 2012.

[59] William McLaren et al. “The Ensembl Variant Effect Predictor”. en. In: Genome biology 17.1 (June 2016), p. 122. issn: 1465-6906. doi: 10.1186/s13059-016-0974-4. url: http://dx.doi.org/10.1186/s13059-016-0974-4.

[60] Brent S Pedersen, Ryan M Layer, and Aaron R Quinlan. “Vcfanno: fast, flexible annotation of genetic variants”. en. In: Genome Biol. 17.1 (Jan. 2016), p. 118.

[61] Johan T den Dunnen et al. “HGVS Recommendations for the Description of Sequence Variants: 2016 Update”. en. In: Hum. Mutat. 37.6 (June 2016), pp. 564–569.

[62] Jennifer L Yen et al. “A variant by any name: quantifying annotation discordance across tools and clinical databases”. en. In: Genome Med. 9.1 (26 01 2017), p. 7.

[63] Todd C Knepper et al. “Key Lessons Learned from Moffitt’s Molecular Tumor Board: The Clinical Genomics Action Committee Experience”. en. In: Oncologist 22.2 (Feb. 2017), pp. 144–151.

[64] Himisha Beltran et al. “Whole-Exome Sequencing of Metastatic Cancer and Biomarkers of Treatment Response”. en. In: JAMA Oncol 1.4 (July 2015), pp. 466–474.

[65] Arezou A Ghazani et al. “Assigning clinical meaning to somatic and germ-line whole-exome sequencing data in a prospective cancer precision medicine study”. en. In: Genet. Med. (26 01 2017).

[66] Alan H Bryce et al. “Experience with precision genomics and tumor board, indicates frequent target identi cation, but barriers to delivery”. en. In: Oncotarget 8.16 (18 04 2017), pp. 27145–27154.

[67] Rong-Fu Wang and Helen Y Wang. “Immune targets and neoantigens for cancer immunotherapy and precision medicine”. en. In: Cell research 27.1 (Jan. 2017), pp. 11–37. issn: 1001-0602, 1748-7838. doi: 10.1038/cr.2016.155. url: http://dx.doi.org/10.1038/cr.2016.155.

[68] Mark Yarchoan et al. “Targeting neoantigens to augment antitumour immunity”. en. In: Nature reviews. Cancer 17.4 (Apr. 2017), pp. 209–222. issn: 1474-175X, 1474-1768. doi: 10.1038/ nrc.2016.154. url: http://dx.doi.org/10.1038/nrc.2016.154.

[69] S T Sherry et al. “dbSNP: the NCBI database of genetic variation”. en. In: Nucleic Acids Res. 29.1 (Jan. 2001), pp. 308–311.

[70] Ethan Cerami et al. “The cBio cancer genomics portal: an open platform for exploring multidimensional cancer genomics data”. en. In: Cancer Discov. 2.5 (May 2012), pp. 401–404.

[71] Jianjiong Gao et al. “Integrative analysis of complex cancer genomics and clinical pro les using the cBioPortal”. en. In: Sci. Signal. 6.269 (Feb. 2013), p. 1.

[72] Carlota Rubio-Perez et al. “Abstract 2983: In silico prescription of anticancer drugs to cohorts of 28 tumor types reveals novel targeting opportunities”. en. In: Cancer Res. 75.15 Supplement (Jan. 2015), pp. 2983–2983.

[73] Abel Gonzalez-Perez et al. “IntOGen-mutations identifies cancer drivers across tumor types”. en. In: Nat. Methods 10.11 (Nov. 2013), pp. 1081–1082.

[74] James R Downing et al. “The Pediatric Cancer Genome Project”. en. In: Nat. Genet. 44.6 (29 05 2012), pp. 619–622.

[75] Jiwen Xin et al. “High-performance web services for querying gene and variant annotation”. en. In: Genome biology 17.1 (June 2016), p. 91. issn: 1465-6906. doi: 10.1186/s13059-016-0953-9. url: http://dx.doi.org/10.1186/s13059-016-0953-9.

[76] Jana Marie Schwarz et al. “MutationTaster2: mutation prediction for the deep-sequencing age”. en. In: Nat. Methods 11.4 (Apr. 2014), pp. 361–362.

[77] Yongwook Choi et al. “Predicting the functional effect of amino acid substitutions and indels”. en. In: PLoS One 7.10 (Aug. 2012), e46688.

[78] Martin Kircher et al. “A general framework for estimating the relative pathogenicity of human genetic variants”. en. In: Nat. Genet. 46.3 (Mar. 2014), pp. 310–315.

[79] Eugene V Davydov et al. “Identifying a high fraction of the human genome to be under selective constraint using GERP++”. en. In: PLoS Comput. Biol. 6.12 (Feb. 2010), e1001025.

[80] Adam Siepel et al. “Evolutionarily conserved elements in vertebrate, insect, worm, and yeast genomes”. en. In: Genome Res. 15.8 (Aug. 2005), pp. 1034–1050.

[81] Katherine S Pollard et al. “Detection of nonneutral substitution rates on mammalian phylogenies”. en. In: Genome Res. 20.1 (Jan. 2010), pp. 110–121.

[82] Lei Bao, Mi Zhou, and Yan Cui. “nsSNPAnalyzer: identifying disease-associated nonsynonymous single nucleotide polymorphisms”. en. In: Nucleic Acids Res. 33. Web Server issue (Jan. 2005), W480–2.

[83] Remo Calabrese et al. “Functional annotations improve the predictive score of human disease-related mutations in proteins”. en. In: Hum. Mutat. 30.8 (Aug. 2009), pp. 1237–1244.

[84] Maximilian Hecht, Yana Bromberg, and Burkhard Rost. “Better prediction of functional effects for sequence variants”. en. In: BMC Genomics 16 Suppl 8 (18 06 2015), S1.

[85] Peng Yue, Eugene Melamud, and John Moult. “SNPs3D: Candidate gene and SNP selection for association studies”. In: BMC Bioinformatics 7.1 (2006), p. 166.

[86] Vikas Pejaver et al. “MutPred2: inferring the molecular and phenotypic impact of amino acid variants”. en. (Sept. 2017).

[87] Majid Masso and Iosif I Vaisman. “AUTO-MUTE: web-based tools for predicting stability changes in proteins due to single amino acid replacements”. en. In: Protein Eng. Des. Sel. 23.8 (Aug. 2010), pp. 683–687.

[88] Paul D Thomas et al. “PANTHER: a library of protein families and subfamilies indexed by function”. en. In: Genome Res. 13.9 (Sept. 2003), pp. 2129–2141.

[89] Alper Uzun et al. “Structure SNP (StSNP): a web server for mapping and modeling nsSNPs on protein structures with linkage to metabolic pathways”. en. In: Nucleic Acids Res. 35. Web Server issue (July 2007), W384–92.

[90] Margarida C Lopes et al. “A combined functional annotation score for non-synonymous variants”. en. In: Hum. Hered. 73.1 (18 01 2012), pp. 47–51.

[91] Yong Mao et al. “CanDrA: cancer-specific driver missense mutation annotation with optimized features”. en. In: PLoS One 8.10 (30 10 2013), e77945.

[92] Christine M Micheel, Christine M Lovly, and Mia A Levy. “My Cancer Genome”. In: Cancer Genet. 207.6 (Jan. 2014), p. 289.

[93] Micheal Hewett et al. “PharmGKB: the Pharmacogenetics Knowledge Base”. en. In: Nucleic Acids Res. 30.1 (Jan. 2002), pp. 163–165.

[94] Linda Huang et al. “The Precision Medicine Knowledge Base: an online application for collaborative editing, maintenance and sharing of structured clinical-grade cancer mutations interpretations”. en. 19 06 2016.

[95] Adrian Tan, Gonalo R Abecasis, and Hyun Min Kang. “Unified representation of genetic variants”. en. In: Bioinformatics 31.13 (Jan. 2015), pp. 2202–2204.

[96] Heng Li et al. “The Sequence Alignment/Map format and SAMtools”. en. In: Bioinformatics 25.16 (15 08 2009), pp. 2078–2079.

[97] Alex H Ramos et al. “Oncotator: cancer variant annotation tool”. en. In: Hum. Mutat. 36.4 (Apr. 2015), E2423–9.

